# Envelope analysis as a tool for identifying epileptic EEG patterns during the sleep-wake cycle in rats

**DOI:** 10.64898/2025.12.27.696681

**Authors:** J. Amaro-Fuenzalida, Javier Díaz, Margarita Bórquez, Carola Mantellero, Patricio Rojas, Alejandro Bassi, Adrián Ocampo-Garcés

**Affiliations:** Universidad de Chile, Facultad de Medicina, Instituto de Ciencias Biomédicas, Laboratorio de Sueño y Cronobiología. Universidad de Chile Facultad de Ciencias Sociales, Departamento de Psicología. Pontificia Universidad Católica de Chile; Universidad de Chile, Facultad de Medicina, Instituto de Ciencias Biomédicas, Laboratorio de Sueño y Cronobiología. International Institute for Integrative Sleep Medicine (WPI-IIIS), University of Tsukuba: Tsukuba; Universidad de Chile, Facultad de Medicina, Instituto de Ciencias Biomédicas, Laboratorio de Sueño y Cronobiología. Universidad de Chile Facultad de Ciencias Sociales, Departamento de Psicología; Universidad de Santiago de Chile, Facultad de Química y Biología, Laboratorio de Neurociencias.; Universidad de Chile, Facultad de Medicina, Instituto de Ciencias Biomédicas, Laboratorio de Sueño y Cronobiología.

**Keywords:** Epilepsy, Sleep, Envelope analysis

## Abstract

**Background:** This research aimed to study electroencephalographic abnormalities in an animal model of temporal lobe epilepsy induced by pilocarpine. By using the coefficient of variation of the envelope (CVE) of electroencephalography (EEG), we characterize and quantify morphological alterations in sleep–wake cycle (SWC) stages.

**Methods:** Epilepsy was induced in 13 rats using a single dose of intraperitoneal pilocarpine injection. A second group of 13 rats served as the control. All the rats underwent polysomnography for at least two days. To do so, three channels were created using four electrodes for EEG and one more for EMG. Channels were built using contralateral lead.

**Results:** Envelope analysis revealed global alterations in EEG morphology. Significant differences were observed in the delta and theta bands between the epileptic and control groups. Epileptic animals showed a near-total suppression of theta rhythm activity during REM sleep. Additionally, we identified a novel pattern of synchronized slow waves (sDelta), distinct from the physiological delta waves observed in non-REM sleep.

**Conclusion:** Identifying both subtle and overt morphological abnormalities in EEG is challenging for human experts and computational methods. By applying CVE analysis to the well-known pilocarpine model, we reveal abnormal EEG dynamics with exceptional summarizing capabilities. For example, a 24-hour EEG can be synthesized into a single, easily interpretable visualization of EEG morphology. This technique and its underlying framework may serve as valuable tools for epilepsy research and potential diagnostic applications.

## 1. Introduction

Epilepsy is a disease characterized by the recurrent and unpredictable appearance of epileptic seizures[1]. In humans, it is usually identified via electroencephalographic (EEG) recordings, which also determine the severity of the disease. In focal epilepsy, EEG can also be used to locate compromised sources[2]. By evaluating electrical brain activity, it has been possible to categorize different types of epilepsy, including temporal lobe epilepsy (TLE), one of the most common types of epilepsy in humans[3]. Epileptic EEG recordings can also show alterations during seizure-free periods, such as interictal activity at different stages of the sleep–wake cycle (SWC), including wakefulness, non-REM sleep, and rapid eye movement (REM) sleep. Each of these stages of the SWC can also be described via EEG recordings[3,4,5].

TLE has a complex relationship with the SWC, as different sleep stages modulate the probability of epileptic seizures[6,7,8,9,10]. Conversely, epileptic seizures can also disorganize the SWC[11,12,13,14]. These alterations have been observed in animal models, such as the pilocarpine model of epilepsy (PME), where a single dose of pilocarpine in rats recreates the main features of TLE, including epileptic seizures. Moreover, other morpohological anomalies such as spikes during interictal periods can be observed during SWC recordings[12].

Most electrophysiological studies on PME have focused on spectral analysis via EEG[15] or have described qualitative alterations during seizure-free periods through visual inspection[12]. While an exhaustive evaluation of the electrophysiological features using both spectral and morphological information is not commonly employed. This approach would allow for the characterization and quantification of abnormalities in the PME.

Fourier-based analysis cannot provide a detailed account of the rich features of epileptic EEG traits in the time domain, as it typically considers the frequency or phase but not the shape of the signal, which is typically evaluated through visual inspection. Therefore, envelope analysis of EEGs has been used to distinguish morphological patterns in the time domain. Specifically, the coefficient of variation of the envelope (CVE) allows for a novel characterization of wild-type SWCs, offering, in addition, a theoretical framework from which the underlying neuronal dynamics can be inferred[16].

The present study focused on the application of CVEs to characterize electroencephalographic abnormalities in an animal model of temporal lobe epilepsy induced by pilocarpine across all stages of the sleep–wake cycle. To achieve this, a canonical description of the sleep–wake cycle is first performed. This characterization will be performed in three steps: (1) qualitatively, through visual inspection; (2) quantitatively, by evaluating the temporal distribution of sleep–wake cycle (SWC) stages; and (3) by assessing the spectral properties of wakefulness, non-REM sleep, and REM sleep. To further enhance this description, the CVE[16] is used to evaluate the morphological properties of the spectrum for each SWC stage. Through this new method, we observed the suppression of certain features commonly associated with REM sleep, while new morphological features emerged that were related to non-REM sleep.

## 2. Materials and methods

### 2.1 Animals

Twenty-six adult male Sprague–Dawley rats (6–7 weeks old, 250–300 g) from a larger doctoral research cohort were used (Mantellero[17]). Thirteen underwent epilepsy induction via a single intraperitoneal pilocarpine injection (PILO group), while thirteen untreated rats served as controls (Control group). All animals were housed individually under a 12:12h light–dark cycle (21–24°C) with food and water ad libitum.

### 2.2 Epilepsy induction

Status epilepticus (SE) was induced in 13 two-week-old rats using the pilocarpine model. Methyl-scopolamine (1mg/kg) was administered 30 min before a single intraperitoneal pilocarpine injection (350mg/kg). Forty-five minutes after SE onset, seizure activity was terminated with diazepam (20mg/kg), following Cavalheiro[18]. Seizures were confirmed using the Racine scale[19]. Control group underwent the same protocol but received saline.

### 2.3 Surgery and polysomnographic recording

Seven days after SE induction, PILO (350 mg/kg) and Control (saline) rats underwent electrode implantation under ketamine/xylazine anesthesia (50/10 mg/kg, i.p.) (see Supplement 1). Using stereotaxic equipment, the skull was drilled at seven sites: bilateral temporal cortex (AP−5.6 mm, ML±4.5 mm), two midline holes (AP−2 mm from lambda, ML+1.5/−0.5mm) for contralateral reference, and three for anchor screws. Electrodes were fixed with dental acrylic, and EMG electrodes were implanted in neck muscles. Postoperative care included antibiotics and analgesics (Supplement 1–2). Spectral analyses used the channel with best signal quality.

Seven days post-surgery (≈ 2 weeks after SE), rats were housed individually in 30×30×25cm cages within 80×80×80cm sound-attenuated, temperature-controlled chambers (21–24 °C) under a 12:12 h light–dark cycle (500 lux, ZT0=07:00). Recordings were made via flexible 40cm cables connected to commutators (PlasticsOne®). EEG(0.3–30Hz) and EMG(30–100Hz) signals were amplified (2000×/5000×), digitized (12bit, 250Hz/channel, Grass15KS; Astro-Med), and stored for offline analysis[20].

All channels were analogically filtered (EEG:0.3–30Hz; EMG:30–100Hz) and amplified (2000×for EEG and 5000×for EMG), digitized (at 12bits, 250Hz per channel, using Grass Model 15KS, Astro-Med, Inc., West Warwick, RI amplifier), and streamed to digital storage for offline analysis[20].

After a 24-hours acclimatization period, rats underwent ≥2 days of undisturbed recording. EEG/EMG data were analyzed in 20s epochs using IgorProV6.1 custom platform. States were scored by two independent observers: wake (low-amplitude EEG + EMG), non-REM (1–4Hz, spindles 10–15Hz, reduced tone), and REM (6–9Hz theta, atonia)[20]. Seizures were verified with an infrared HD-CMOS camera. To quantify sleep–wake cycle (SWC) distribution, each 24-hour period was divided into eight 3-hour octants, and the number of epochs per stage and octant was used to calculate time spent in each state.

### 2.4 Data analysis and statistics

To characterize EEG alterations CVE was calculated[16, 21]. EEG epochs (20s, 50 % overlap) were filtered into delta (0.5–4Hz), theta (4–10Hz), and sigma (11–16Hz) bands. For each epoch, we computed the coefficient of variation of the envelope (CVE)[21] and root mean square (RMS)[22]. Gaussianity fingerprint 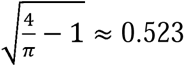 (±1.96 standard deviations) was used to classify morphology as sine-like, Gaussian-like, or pulse-like. Epochs were mapped in the Envelope Characterization Space (ECS)[16], a CVE×RMS scatterplot generated for each rat and frequency band (Figure 1). The x-axis (CVE) reflects morphology relative to the 0.523 threshold and its confidence intervals[16], while the y-axis (RMS) uses the median to separate high vs. low amplitude. The ECS was divided into four quadrants (CVE below/above 0.523×RMS below/above median), each representing distinct oscillatory patterns. Figure 1 illustrates representative epochs across the CVE spectrum (low, mid, high) and the quadrant layout.

**Figure 1.**
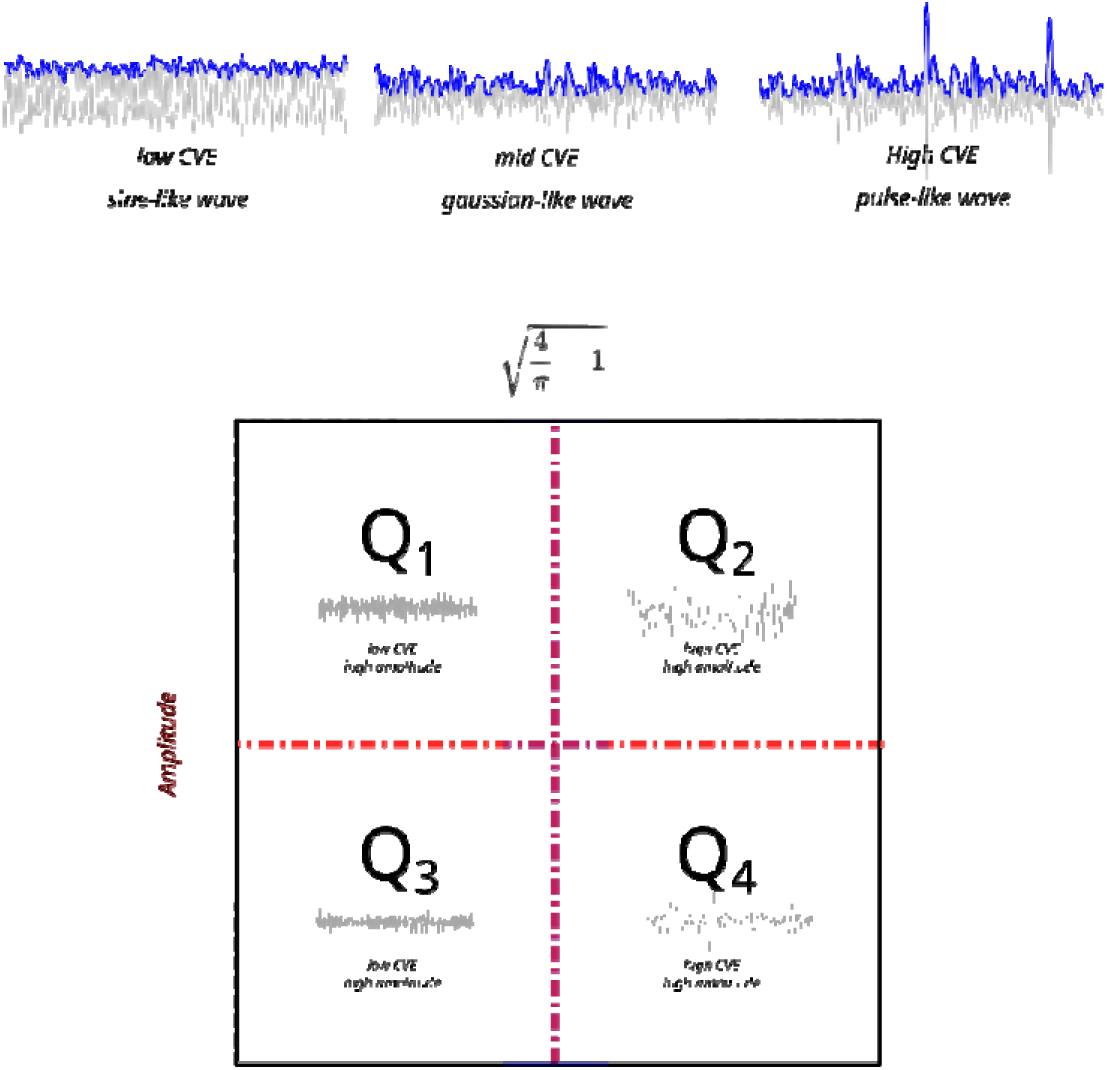
Envelope analysis description and Envelope Characterization Space. Top: A low CVE value describes a sine-like envelope, a medium CVE value generates a Gaussian-like envelope, and a high CVE value produces a pulse-like envelope. To perform the CVE analysis, all recording periods were evaluated according to the sampling distribution of CVE in each band (delta, theta, sigma). Subsequently, it was determined whether the periods belonged to the sampling distribution of Gaussianity or not. This resulted in two rejection zones: one significantly lower than Gaussian noise (low CVE, sine-like wave) and another significantly higher than noise (high CVE, pulse-like wave). Bottom: Envelope Characterization Space[16]. After CVE calculation and classification, we use the amplitude of the epochs to provide a position in Envelope Characteristics Space. Additionally, median amplitude and CVE gaussianity fingerprint is used to create four quadrants. Q1: low CVE with high amplitude. Q2: high CVE with high aplitude, Q3: low CVE with low amplitude, Q4: high CVE with low amplitude. Each quadrant includes a representative example epoch located on each position of envelop characterization space.

For each rat and band, we calculated epoch counts per quadrant across all recording days. Statistical analyses included repeated-measures ANOVA with Greenhouse–Geisser correction and Bonferroni post hoc tests; effect sizes are reported as partial η² and Cohen’s d[23].

## 3. Results

Figure 2A shows a representative epileptic seizure. Spontaneous seizures were observed in all pilocarpine-treated rats during induction, and during EEG sessions seizures were detected in 10/13 animals, averaging 1.8±2.6 seizures over the recording period. While 1.3(SD=2.4) seizures occurred during the light phase, only 0.5(SD=0.9) was observed during the dark phase (Figure 2B). More seizures occurred during the light (1.3±2.4) than dark phase (0.5 ± 0.9) (paired t=2.47, p=0.01, d=0.46; Figure 2B). Hourly distributions (Figure 2C) revealed sporadic interictal spikes interrupting otherwise normal activity, consistent with previous reports[12,15,24].

**Figure 2.**
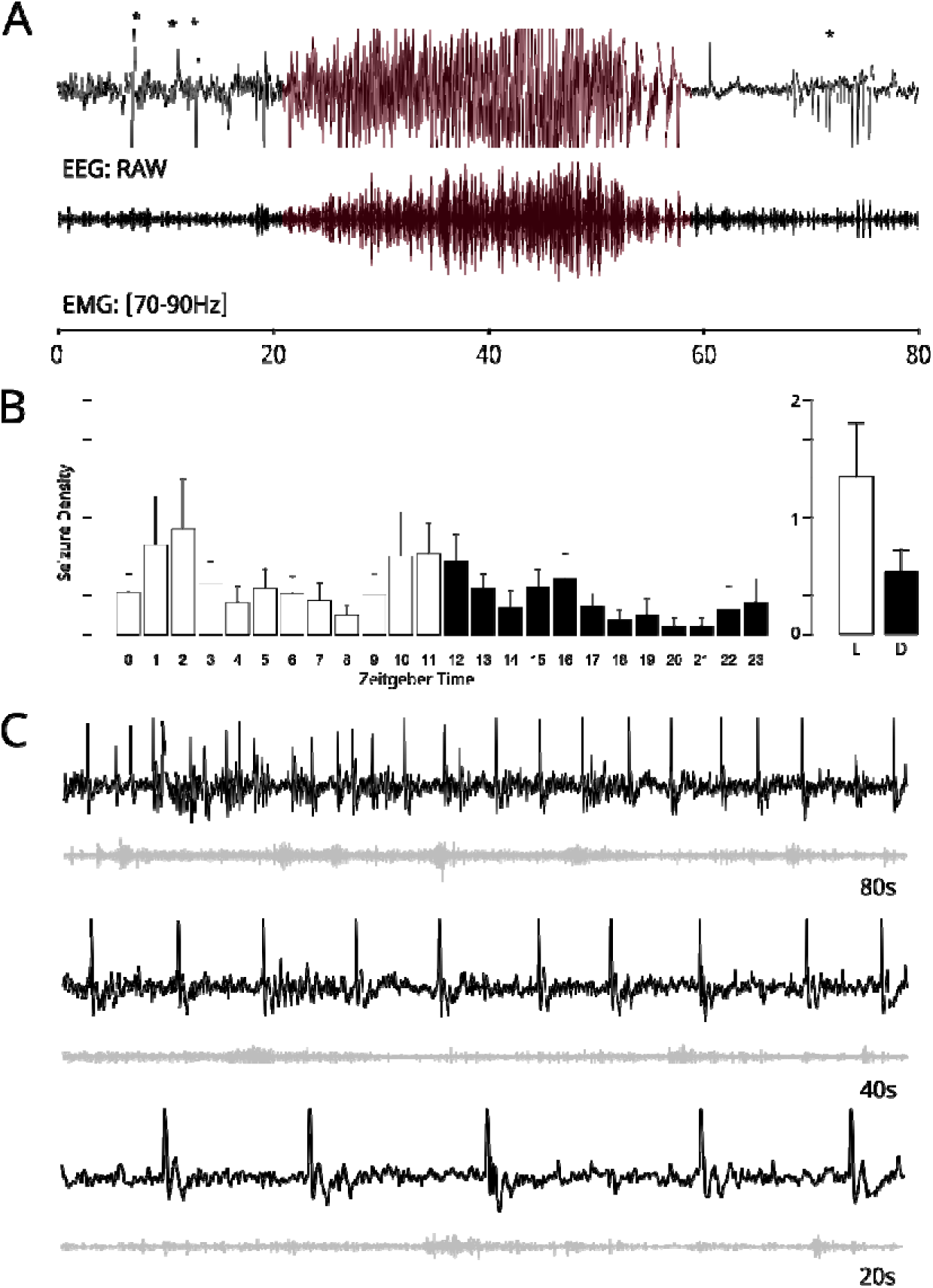
EEG activity of epileptic activity in a representative subject. A: Example of an epileptic seizure. The upper panel shows EEG activity, whereas the lower panel represents EMG activity. (*) Indicates preictal and post ictal patterns. B: Temporal distribution of seizures relative to Zeitgeber Time (ZT). ZT0 corresponds to lights-on (07:00 AM) and ZT12 to lights-off (07:00 PM). The x-axis represents circadian hours from ZT0 to ZT23. White bars (L) indicate the light phase (ZT0–ZT11), and black bars (D) indicate the dark phase (ZT12–ZT23). C: Example of interictal spike activity during wakefulness in an epileptic animal. The same EEG segment is displayed at three temporal resolutions (80, 40, and 20 s) to highlight morphological features. Gray traces indicate EMG activity.

Control rats exhibited canonical EEG patterns for wakefulness, NON-REM, and REM sleep[5, 25], whereas PILO rats showed disrupted morphology across all states, including an almost complete loss of theta rhythm during REM and reduced amplitude (Figure 3, left). NON-REM patterns were visually similar between groups. Spectral analyses (Figure 3, right) confirmed a 4–10Hz theta peak in controls during wakefulness and REM, which was reduced in PILO rats. Theta amplitude during REM was significantly higher in controls than PILO (t=5.84, p=0.01, d=1.7), but no group difference was found during wakefulness (t=0.19, p=0.42). Within controls, theta amplitude was greater in REM than wakefulness (paired t=4.65, p < 0.001, d=0.9), whereas PILO rats showed no difference (t=−1.51, p=0.92).

**Figure 3.**
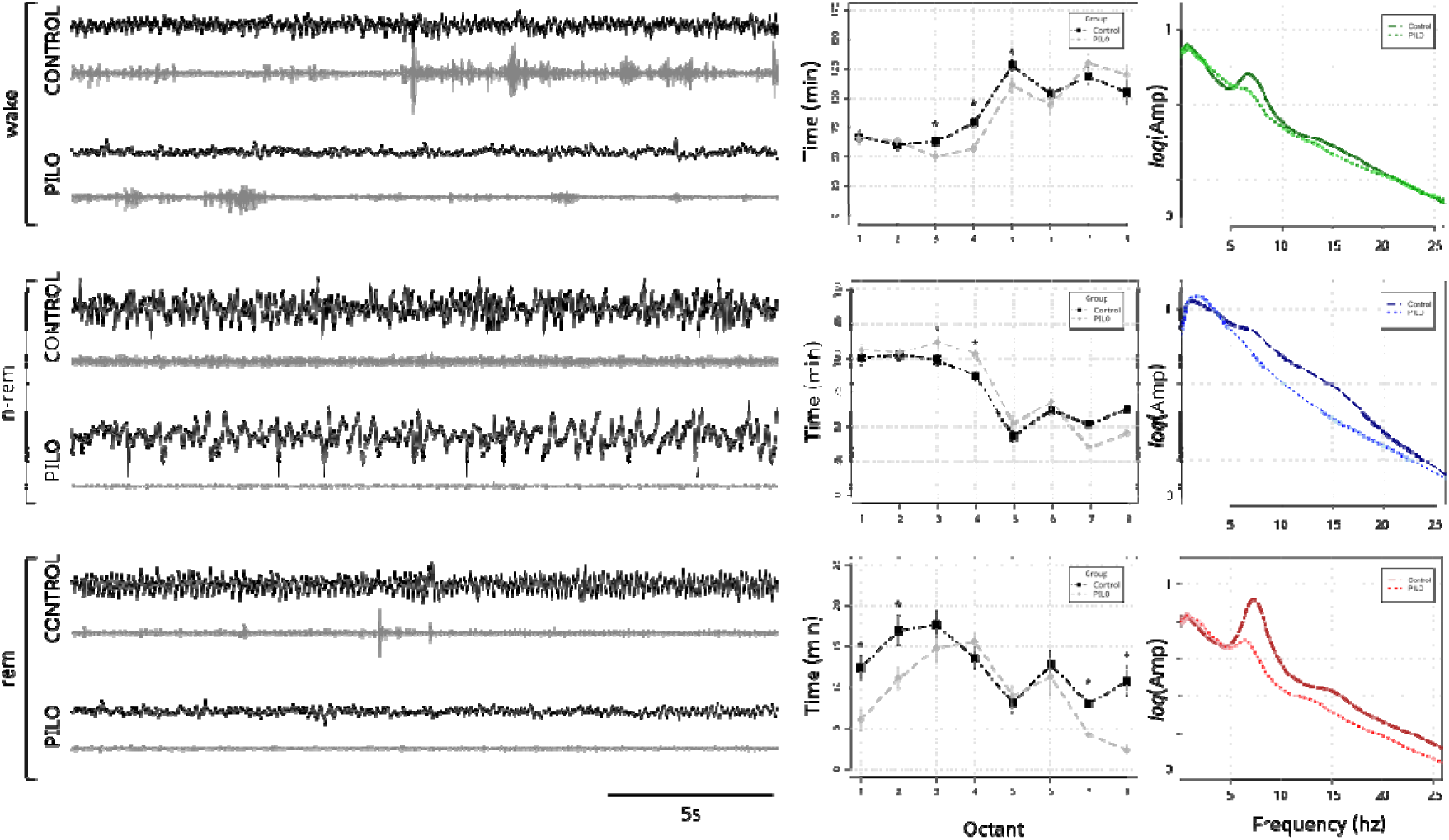
EEG activity and sleep–wake cycle activity. Left: The three stages of the sleep-wake cycle for Control and PILO representative subjects are presented. Middle: Time distribution of stages of the sleep-wake cycle in the control and PILO groups. The (*) indicates significant differences in the amount of time invested in each stage, during each octant. Control group describe significant more time on wakefulness in octant three, four and five. While the same happened with rem sleep, in first, second, seventh and eighth octant. Epileptic rats, invest significant more time in n-rem during octant three and four. Right: Spectral analysis of EEGs for the control and PILO groups as a function of the sleep-wake cycle stage. There are no significant differences in theta band (4-10Hz) in PILO group when rem and wakefulness are compared. While this difference can be observed in Control group.

Analysis of 24h SWC distributions (averaged across both recording days) showed that both groups spent more time awake during the dark phase and asleep (NON-REM and REM) during the light phase, with REM increasing during the second half of the light period in all animals (Figure 3, middle). For wakefulness, there were significant main effects of octant (F(4.4,107.2)=50.10, p<0.001) and group × octant interaction (F(4.4,107.2)=2.78, p=0.025). Controls spent more time awake than PILO rats during the third (t=2.25, p=0.034, d=0.88), fourth (t=4.47, p=0.01, d=1.75), and fifth (t=2.31, p=0.03, d=0.9) octants.

For non-REM sleep, there were significant octant effects (F(4,103)=67.95, p<0.001) and group×octant interaction (F(4,103)=3.83, p=0.005). PILO rats spent more time in NON-REM than controls during the third (t=−2.43, p=0.02, d=0.95) and fourth (t=−3.97, p=0.01, d=1.56) octants.

For REM sleep, both groups showed significant octant effects (F(5,117)=17.78, p<0.001) and group × octant interaction (F(5,117)=3.89, p=0.005). Controls spent more time in REM than PILO rats during the first (t=3.39, p=0.002, d=1.33), second (t=2.59, p=0.016, d=1.01), seventh (t=2.31, p=0.03, d=0.9), and eighth (t=4.42, p=0.01, d=1.73) octants.

CVE analysis was applied to quantify EEG morphology (Figure 4). In controls, low-amplitude epochs clustered around CVE≈0.523 across bands, reflecting Gaussian-like wakefulness. In delta, high-CVE/high-amplitude epochs corresponded to NON-REM, while lower amplitudes near 0.523 aligned with REM and wake. In theta, high amplitude showed both high CVE (NON-REM) and low CVE (REM) clusters, the latter indicating sinusoidal theta during REM. Sigma displayed a pattern similar to delta. In PILO rats, low amplitudes also corresponded to Gaussian-like wakefulness, but delta-band epochs of median amplitude showed high CVE, indicating interictal spikes. With increasing amplitude, CVE trended toward 0.523 or below, suggesting rhythmicity. In theta, the high-CVE cluster remained but the low-CVE REM cluster was absent, matching the loss of REM theta noted earlier. Sigma clusters were similar between groups.

**Figure 4.**
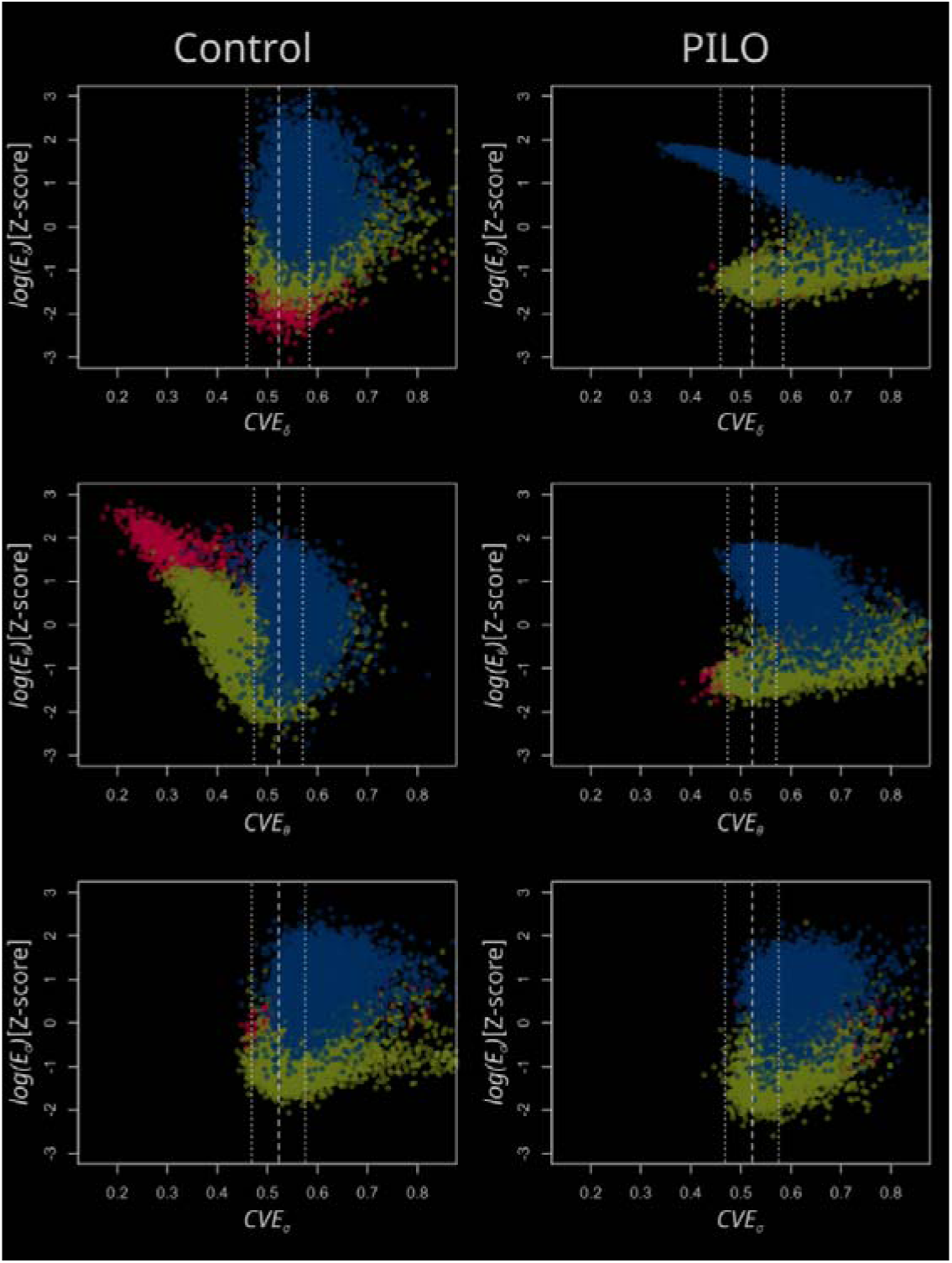
Envelope characterization space. Envelope Characterization Space (ECS) for a representative subject from each group (columns). Each row corresponds to a different frequency band (Delta, Theta, and Sigma). The Y-axis represents each band’s standardized power, whereas the X-axis represents the CVE values. Each dot represents an epoch categorized by sleep stage: blue represents non-REM sleep, red represents REM sleep, and green represents wakefulness. Supplement 3 shows the ECS for all PILO rats, while the control ECS is summarized in Diaz et al., 2018[16].

Significant group differences were found in the first ECS quadrant (low amplitude, low CVE) for delta and theta bands. PILO rats spent more time in the delta first quadrant than controls (t□□.□=−5.8, p<0.001, d=−2.28), whereas controls spent more time in theta (t□□.□=2.26, p<0.05, d=0.89). No differences were observed in sigma (t□□.□=0.99, p=0.45). PILO rats exhibited a striking aberrant pattern in the delta band, characterized by an accumulation of low-CVE, high-delta epochs in the first ECS quadrant, indicating sine-like delta oscillations rather than the pulse-like delta activity typical of controls[16]. We termed this distinctive pattern sinusoidal delta (sDelta). sDelta represents high-amplitude, low-CVE activity, reflecting a flatter envelope than that normally observed in physiological NON-REM sleep. PILO rats spent significantly more time in sDelta than controls (t□□=5.97, p<0.001, d=1.71; Figure 5). Based on amplitude criteria and EMG RMS values below the 25th percentile, sDelta was classified as occurring during resting periods and were classified as NON-REM by visual inspection.

**Figure 5.**
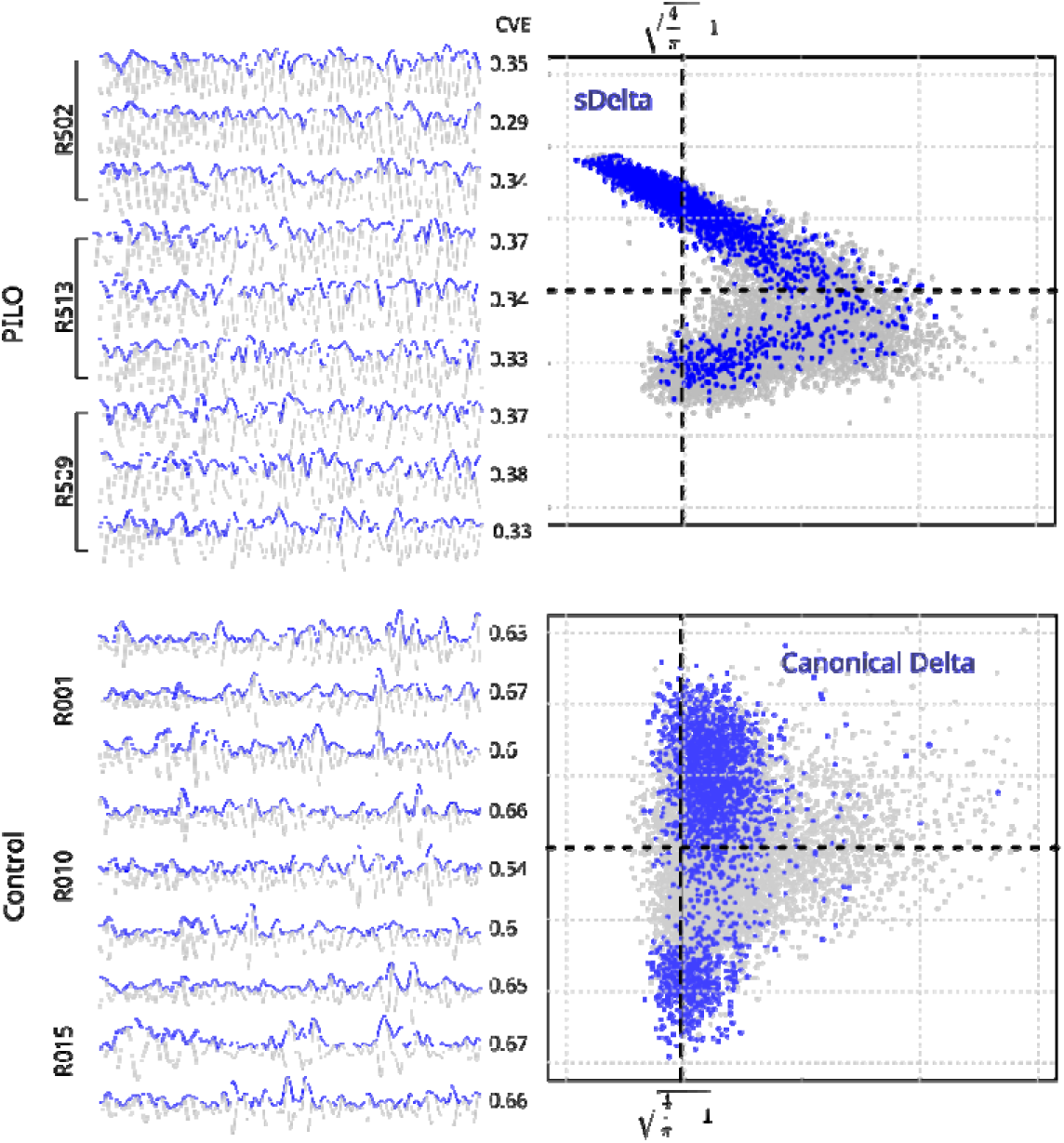
Envelope characterization space and EMG activity. Nine examples of epochs with the highest amplitude in the delta band (0.5–4Hz) for each PILO and control group. Each sample includes a CVE value. Importantly, for the PILO group, the CVE is low (sine-like wave), whereas the control group has values greater than –0.523, which indicates pulse-like activity. In the right panel, the ECVs for PILO and Control are plotted. The x-axis represents the CVE, whereas the y-axis represents the amplitude of the signal. The gray dots represent all the epochs, whereas the blue dots indicate all the epochs with EMG amplitudes below P25 (resting). Importantly, both the PILO group and the control group show two clusters, whereas low-amplitude clusters are similar in both groups, with Gaussian-like values, when the amplitude increases, the blue dots in the PILO group reach the first quadrant in the ECV. Most of the epochs in the control group belong to the second quadrant of the ECV. Representative waveform examples for each ECS quadrant (including the novel sDelta pattern) are provided in Figure 1

## 4. Discussion

### EEG features

Traditionally, EEG pattern identification for physiological or pathological states relies on visual inspection. Using this approach, we found that seizures predominated during the light phase (Figure 1B), suggesting circadian modulation of seizure activity, consistent with human temporal lobe epilepsy studies linking seizures to specific sleep states and circadian transitions[6,7,8], potentially mediated by hypothalamic and thalamocortical rhythms. Visual inspection showed reduced wakefulness and increased non-REM sleep in PILO rats, indicating homeostatic sleep reorganization. Hence, frequency-based analysis revealed more amplitude on delta NON-REM and suppressed theta activity during REM compared to controls.

This theta suppression may stem from pilocarpine’s effects, which disrupt cholinergic signaling. Because REM theta depends on muscarinic activation, sustained overstimulation or neurotoxicity could impair its generation or propagation. This aligns with the hippocampus’s role as a REM theta pacemaker and how its disruption can facilitate seizures in other models[26]. REM sleep may exert a protective effect against seizures, and pharmacological REM induction (e.g., carbachol) has been linked to increased seizure susceptibility. Thus, the loss of REM-related theta in the PILO model may reflect both hippocampal damage and the loss of a protective mechanism against epileptiform activity[7]. Moreover, recurrent interictal discharges in temporal lobe epilepsy may disrupt the synchronization required for theta expression.

In PILO rats, REM suppression made REM and wakefulness visually indistinguishable due to their similar EEG patterns. Since amplitude-based spectral analysis misses key morphological features, complementary methods are needed to characterize EEG signals accurately[27,28,29].

### CVE features

CVE analysis provides a quantitative framework for characterizing EEG morphology. Gaussian activity typically exhibits 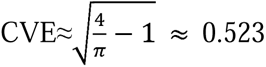, close to Rayleigh distribution coefficient. Thresholds around this value, defined statistically, allow the objective classification of rhythmic (low CVE), Gaussian (≈0.523), and phasic (high CVE) patterns. These categories, however, should not be interpreted rigidly; rather, CVE should be viewed as a continuous metric, capturing the gradual nature of EEG morphology—lower values reflect more sinusoidal activity, whereas higher values indicate transient, nonstationary events[16]. Although the 0.523 threshold originates from theoretical models, CVE analysis has also been applied successfully to human EEG[30], suggesting that deviations from Gaussianity can be informative for identifying pathological transitions, such as those observed in epilepsy.

The application of CVE analysis in the PME model demonstrates its utility for quantitatively characterizing EEG alterations. Specifically, the envelope characterization space (CVE vs. log(mean(env))) revealed distinct clustering patterns between control and pilocarpine-treated rats (Figure 4). Furthermore, abnormal EEG patterns identified by CVE exhibited both circadian and brain state specificity (Figures 1A, C, and 3). Particularly notable is the circadian organization of synchronized delta oscillations (sDelta), a hallmark feature of the PME phenotype.

Typically, a CVE-based generative model accounts for delta wave heterogeneity by modeling them as transient events at different rates: low rates yield phasic bursts with high CVE, whereas high rates produce Gaussian-like activity (CVE ≈ 0.523)[16]. In the PME model, delta-band CVE and amplitude show an inverse linear correlation, especially during high-amplitude epochs (Figure 4). Uniquely, PME exhibits high-amplitude epochs with abnormally low CVE, indicating pathological rhythmic delta activity. This CVE-detected delta alteration is functionally relevant, as waveform morphology may reveal mechanisms such as sDelta, potentially linked to epilepsy-related functional disturbances[28].

PME is a valuable model for TLE, reproducing key features such as hippocampal and amygdalar sclerosis, seizure activity, interictal discharges, and propagation patterns [31,32,33]. though it has been criticized for variability linked to age, sex, and strain [34]. We hypothesized that hippocampal sclerosis would still shape brain oscillations, and CVE analysis confirmed a consistent pattern of EEG morphological alterations across subjects. As shown in Supplement 3, CVE-based characterization exhibited limited inter-subject variability, indicating that PME induces robust EEG alterations detectable by CVE, regardless of seizure frequency or duration. Future work should examine whether individual differences in ECS geometry (e.g., cluster density, spread, slope) correlate with clinical markers of epilepsy severity, to assess the predictive or diagnostic potential of CVE metrics.

### What is sDelta?

sDelta appears mainly during resting periods with low EMG activity, suggesting a link to physiological sleep states. It emerges predominantly under high sleep pressure, but its distinct morphology and absence in controls indicate it is not simply a variant of NON-REM or REM sleep. Considering this, the key question is what drives the transition from Gaussian-like to sine-like activity in PMEs. One explanation involves thalamo-cortical networks, whose alterations may impose abnormal rhythms on cortical activity[34,35,36], as seen in spike-and-wave discharges (SWDs)[24]. Alternatively, changes in cortical excitability and propagation through parallel pathways may underlie this pattern[37]. Morphological changes identified via CVE raise the question of whether all NON-REM epochs are truly physiological. sDelta (Figure 5) might reflect a morphological alteration of NON-REM, produced by PME-induced histological anomalies[33,34,38] affecting slow-wave generators, leading to pulsatile or sinusoidal order. Testing this requires examining sDelta’s temporal distribution under different levels of sleep pressure. If sDelta reflects altered NON-REM, it should increase during high sleep pressure, regulated by the sleep S process[39,40,41].

sDelta could also involve hippocampal damage caused by pilocarpine and REM disruptions[31,32,33,34]. In rodents, REM sleep is marked by a hippocampal theta rhythm, forming a sine-like 4–10Hz pattern[16]. Like delta, sDelta is sinusoidal but slower (0.5–4Hz), possibly reflecting a slowed theta oscillator due to hippocampal damage[33,35,35], rather than altered NON-REM. Since pilocarpine can also modulate cortical–hippocampal connectivity during REM[42], this explanation could be plausible.

Since sDelta exhibits unique EEG features, it overlaps with NON-REM (amplitude) and REM (morphology), future studies should assess whether sDelta has restorative functions akin to deep sleep[43] or reflects epilepsy-related cortical disconnection[37]. Homeostatic regulation aligning with the synaptic homeostasis hypothesis[43], which posits that slow waves and spindles restore synaptic efficacy after waking. The increased NON-REM time in PILO rats may reflect compensation for reduced sleep efficiency, and quantifying sDelta could offer a new diagnostic window into sleep alterations in refractory epilepsy, especially when standard EEG misses such changes.

### CVE advantages and disadvantages

CVE quantifies EEG morphology and, unlike spectral analyses focused on power, captures envelope variability over time, highlighting physiologically relevant waveform differences often overlooked[27,28,29,44]. While a signal’s spectral profile remains unchanged after phase randomization, its CVE can vary substantially. Hence, CVE is conceptually simple, computationally efficient, requires no event detection, and applies uniformly across continuous EEG. In our study, CVE revealed systematic deviations in epileptic rats not detected by visual inspection or spectral power, aligning with findings by Truccolo et al.[45], who observed persistent neuronal firing changes between seizures. This suggests that CVE could potentially detect network-level disturbances, identifying abnormal states even without clinical events. Nevertheless, further work is needed to test whether CVE correlates with human clinical markers.

Despite these advantages, CVE has limitations. It is calculated per channel, lacking spatial resolution unless complemented by other methods. Its values are sensitive to filter bandwidth and window length, requiring strict standardization for comparisons[16,47]. Interpretation may be challenging, while CVE identifies deviations from Gaussian-like envelope behavior, it does not specify the source or nature of the deviation. For example, a high CVE could reflect sharp pathological activity, transient artifacts, or sparse events [46]. Because CVE distributions under non-Gaussian conditions lack closed forms, empirical thresholds must be derived from Gaussian noise models, introducing dependence on surrogate benchmarks. CVE is also less established than techniques like spectral entropy[4] or 1/f slope analysis[48].

Although applications in epilepsy are limited, CVE has been used to examine slow-wave stability[49], age-related sleep changes[50], and Gaussian deviations in oscillations[30] in humans. These studies highlight its potential to provide fine-grained, physiologically meaningful metrics. Overall, CVE offers a valuable complement to existing EEG analyses, but its parameter interpretation calls for cautious use and methodological refinement.

## 5. Declarations

### Ethics approval and consent to participate

The animals in the experimental (PILO) group were recorded as part of a doctoral research project conducted at the Universidad de Santiago de Chile (Mantellero Gutiérrez, 2017). While the original protocol number is not publicly available, all procedures adhered to institutional and national ethical guidelines for animal research and were conducted under academic and ethical supervision[17]. Electroencephalographic procedures and housing conditions were consistent with protocols approved by the Bioethics Committee of the Faculty of Medicine, Universidad de Chile, under approval number CBA 0797 FMUCH.

### Consent for publication

Not applicable.

### Authors’ contributions

- Margarita Bórquez, Carola Mantellero, and J. Amaro-Fuenzalida performed the experimental procedures for polysomnography and epilepsy induction.
- J. Amaro-Fuenzalida conducted the data analysis and signal processing under the supervision of Javier Díaz.
- Alejandro Bassi supervised the development of recording devices and software.
- Adrián Ocampo-Garcés and Margarita Bórquez advised J. Amaro-Fuenzalida on behavioral and sleep-wake cycle analysis.
- Patricio Rojas advised Carola Mantellero and provided expertise on epilepsy management.
- All recordings were performed in Adrián Ocampo-Garcés’ laboratory using devices developed by Alejandro Bassi.

### Availability of data and materials

The datasets used and/or analysed during the current study are available from the corresponding author on reasonable request.

### Figures

All figures have been submitted as separate high-resolution files in PDF format, following journal requirements. Figure legends are included within the manuscript.

## References

[1] Fisher RS, Acevedo C, Arzimanoglou A, Bogacz A, Cross JH, Elger CE, Engel J, Jr., Forsgren L, French JA, Glynn M, Hesdorffer DC, Lee BI, Mathern GW, Moshe SL, Perucca E, Scheffer IE, Tomson T, Watanabe M, Wiebe S. ILAE official report: a practical clinical definition of epilepsy. Epilepsia 2014;55: 475–82.10.1111/epi.12550.

[2] Sanz-García A, Vega-Zelaya L, Pastor J, Sola RG, Ortega GJ. Donde comienza el periodo postictal en la epilepsia de lobulo temporal? Hacia una definición cuantitativa. Revista de Neurología 2017;64: 337–346.10.33588/rn.6408.2016269.

[3] Wiebe S. Epidemiology of Temporal Lobe Epilepsy. Canadian Journal of Neurological Sciences 2000;27: S6–S10.10.1017/s0317167100000561.

[4] Tellez-Zenteno JF, Hernandez-Ronquillo L. A review of the epidemiology of temporal lobe epilepsy. Epilepsy Res Treat 2012;2012: 630853.10.1155/2012/630853.

[5] Siegel JM. The neurobiology of sleep. Semin Neurol 2009;29: 277–96.10.1055/s-0029-1237118.

[6] Crespel A, Baldy-Moulinier M, Coubes P. The relationship between sleep and epilepsy in frontal and temporal lobe epilepsies: practical and physiopathologic considerations. Epilepsia 1998;39: 150–157.10.1111/j.1528-1157.1998.tb01352.x.

[7] Shouse MN, Farber PR, Staba RJ. Physiological basis: how NON-REM sleep components can promote and REM sleep components can suppress seizure discharge propagation. Clinical Neurophysiology 2000;111: S9–S18.10.1016/s1388-2457(00)00397-7.

[8] Herman ST, Walczak TS, Bazil CW. Distribution of partial seizures during the sleep–wake cycle: differences by seizure onset site. Neurology 2001;56: 1453–1459.10.1212/wnl.56.11.1453.

[9] Kumar P, Raju TR. Seizure susceptibility decreases with enhancement of rapid eye movement sleep. Brain research 2001;922: 299–304.10.1016/S0006-8993(01)03174-2.

[10] Badawy RA, Curatolo JM, Newton M, Berkovic SF, Macdonell RA. Sleep deprivation increases cortical excitability in epilepsy: syndrome-specific effects. Neurology 2006;67: 1018–1022.10.1212/01.wnl.0000237392.64230.f7.

[11] Bazil CW, Castro LH, Walczak TS. Reduction of rapid eye movement sleep by diurnal and nocturnal seizures in temporal lobe epilepsy. Archives of neurology 2000;57: 363–368.10.1001/archneur.57.3.363.

[12] Matos G, Tsai R, Baldo M, de Castro I, Sameshima K, Valle A. The sleep–wake cycle in adult rats following pilocarpine-induced temporal lobe epilepsy. Epilepsy y Behavior 2010;17: 324–331.10.1016/j.yebeh.2009.11.015.

[13] Hofstra W, Gordijn M., van der Palen, J., van Regteren, R., Grootemarsink, B., y, de Weerd A. Timing of temporal and frontal seizures in relation to the circadian phase: a prospective pilot study. Epilepsy research 2011;94: 158–162.10.1016/j.eplepsyres.2011.01.015.

[14] Yi PL, Chen YJ, Lin CT, Chang FC. Occurrence of epilepsy at different zeitgeber times alters sleep homeostasis differently in rats. Sleep 2012;35.10.5665/sleep.2238.

[15] Bortel A, Lévesque M, Biagini G, Gotman J, Avoli M. Convulsive status epilepticus duration as determinant for epileptogenesis and interictal discharge generation in the rat limbic system Neurobiology of disease 2010;40: 478–489. 10.1016/j.nbd.2010.07.015.

[16] Diaz J, Bassi A, Coolen A, Vivaldi EA, Letelier JC. Envelope analysis links oscillatory and arrhythmic EEG activities to two types of neuronal synchronization. Neuroimage 2018;172: 575–585.10.1016/j.neuroimage.2018.01.063.

17. Mantellero Gutiérrez, C. A. (2017). Bumetanida potencia el efecto farmacológico de fenobarbital, a nivel electroencefalográfico y conductual, en un modelo animal de epilepsia del lóbulo temporal. Universidad de Santiago de Chile.

[18] Cavalheiro EA. The pilocarpine model of epilepsy. The Italian Journal of Neurological Sciences 1995;16: 33–37.10.1007/BF02229072.

[19] Luttjohann A, Fabene PF, van Luijtelaar G. A revised Racine’s scale for PTZ-induced seizures in rats. Physiol Behav 2009;98: 579–86.10.1016/j.physbeh.2009.09.005.

[20] Castro-Faundez J, Diaz J, Ocampo-Garces A. Temporal Organization of the Sleep-Wake Cycle under Food Entrainment in the Rat. Sleep 2016;39: 1451–65.10.5665/sleep.5982.

[21] Diaz JA, Arancibia JM, Bassi A, Vivaldi EA. Envelope analysis of the airflow signal to improve polysomnographic assessment of sleep disordered breathing. Sleep 2014;37: 199–208.10.5665/sleep.3338.

[22] Rangayyan R. Biomedical Signal Analysis. Second ed: John Wiley y Sons; 2015.

[23] Lakens D. Calculating and reporting effect sizes to facilitate cumulative science: a practical primer for t tests and ANOVAs. Front Psychol 2013;4: 863.10.3389/fpsyg.2013.00863.

[24] Loddenkemper T, Fernández IS, Peters JM. Continuous spike and waves during sleep and electrical status epilepticus in sleep. Journal of Clinical Neurophysiology 2011;28: 154–164.10.1097/WNP.0b013e31821213eb.

[25] Ocampo-Garcés A, Vivaldi EA. Short-term homeostasis of REM sleep assessed in an intermittent REM sleep deprivation protocol in the rat. Journal of sleep research 2002;11: 81–89.10.1046/j.1365-2869.2002.00281.x.

[26] Sedigh-Sarvestani, M., Thuku, G. I., Sunderam, S., Parkar, A., Weinstein, S. L., Schiff, S. J., & Gluckman, B. J. (2014). Rapid eye movement sleep and hippocampal theta oscillations precede seizure onset in the tetanus toxin model of temporal lobe epilepsy. Journal of Neuroscience, 34(4), 1105–1114.

[27] Cole SR, Voytek B. Brain Oscillations and the Importance of Waveform Shape. Trends Cogn Sci 2017;21: 137–149.10.1016/j.tics.2016.12.008.

[28] Diaz J, Razeto-Barry P, Letelier JC, Caprio J, Bacigalupo J. Amplitude modulation patterns of local field potentials reveal asynchronous neuronal populations. J Neurosci 2007;27: 9238–45.10.1523/JNEUROSCI.4512-06.2007.

[29] Ocak H. Automatic detection of epileptic seizures in EEG using discrete wavelet transform and approximate entropy. Expert Systems with Applications 2009;36: 2027–2036.10.1016/j.eswa.2007.12.065.

[30] Hidalgo, V. M., Díaz, J., Mpodozis, J., & Letelier, J. C. (2022). Envelope analysis of the human alpha rhythm reveals EEG gaussianity. IEEE Transactions on Biomedical Engineering, 70(4), 1242–1251.

[31] Covolan L, Mello LEAM. Temporal profile of neuronal injury following pilocarpine or kainic acid-induced status epilepticus. Epilepsy research 2000;39(2): 133–152.10.1016/s0920-1211(99)00119-9.

[32] Toyoda I, Bower MR, Leyva F, Buckmaster PS. Early activation of ventral hippocampus and subiculum during spontaneous seizures in a rat model of temporal lobe epilepsy. Journal of Neuroscience 2013;33: 11100–11115.10.1523/JNEUROSCI.0472-13.2013.

[33] Polli RS, Malheiros JM, Dos Santos R, Hamani C, Longo BM, Tannus A, Mello LE, Covolan L. Changes in Hippocampal Volume are Correlated with Cell Loss but Not with Seizure Frequency in Two Chronic Models of Temporal Lobe Epilepsy. Front Neurol 2014;5: 111.10.3389/fneur.2014.00111.

[34] Curia G, Longo D, Biagini G, Jones RS, Avoli M. The pilocarpine model of temporal lobe epilepsy. Journal of neuroscience methods 2008;172: 143–157.10.1016/j.jneumeth.2008.04.019.

[35] Steriade M, Timofeev I. Neuronal Plasticity in Thalamocortical Networks during Sleep and Waking Oscillations. Neuron 2003;37: 563–576.10.1016/S0896-6273(03)00065-5.

[36] Hagler DJ, Jr., Ulbert IA-O, Wittner LA-O, Erőss L, Madsen JR, Devinsky OA-O, Doyle WA-O, Fabó DA-O, Cash SS, Halgren EA-O. Heterogeneous Origins of Human Sleep Spindles in Different Cortical Layers. The Journal of neuroscience : the official journal of the Society for Neuroscience 2018;38: 3013–3025.10.1523/JNEUROSCI.2241-17.2018.

[37] Polack PO, Guillemain I, Hu E, Deransart C, Depaulis A, Charpier S. Deep layer somatosensory cortical neurons initiate spike-and-wave discharges in a genetic model of absence seizures. J Neurosci 2007;27: 6590–9.10.1523/JNEUROSCI.0753-07.2007.

[38] Hester MS, Danzer SC. Accumulation of abnormal adult-generated hippocampal granule cells predicts seizure frequency and severity. Epilepsy Behav. 2013;38:13–21. doi: 10.1016/j.yebeh.2013.04.011.

[39] Drinkenburg WHIM, Coenen AML, Vossen JMH, Van Luijtelaar ELJM. Sleep deprivation and spike-wave discharges in epileptic rats. Sleep 1995;18(4): 252–256.10.1093/sleep/18.4.252.

[40] Borbely AA, Daan S, Wirz-Justice A, Deboer T. The two-process model of sleep regulation: a reappraisal. J Sleep Res 2016;25: 131–43.10.1111/jsr.12371.

[41] Achermann P, Dijk DJ, Brunner DP, Borbély AA. A model of Human Sleep Homeostasis Based on EEG Slow-Wave Activity. Brain research bulletin 1993;31(1-2): 97–113.10.1016/0361-9230(93)90016-5.

[42] Tejada S, González JJ, Rial RV, Coenen AM, Gamundí A, Esteban S. Electroencephalogram functional connectivity between rat hippocampus and cortex after pilocarpine treatment. Neuroscience. 2010;165(2):621–31. doi: 10.1016/j.neuroscience.2009.10.031.

[43] Tononi, G., & Cirelli, C. (2005). Sleep function and synaptic homeostasis. Sleep Medicine Reviews, 10(1), 49–62. 10.1016/j.smrv.2005.05.002

[44] Bruns, A. (2004). Fourier-, Hilbert- and wavelet-based signal analysis: Are they really different approaches? Journal of Neuroscience Methods, 137(2), 321–332. 10.1016/j.jneumeth.2004.03.002

[45] Truccolo, W., Donoghue, J. P., Hochberg, L. R., Eskandar, E. N., Madsen, J. R., Anderson, W. S., & Brown, E. N. (2011). Single-neuron dynamics in human focal epilepsy. Nature Neuroscience, 14(5), 635–641. 10.1038/nn.2782

[46] Tatum, W. O. IV (2021). Handbook of EEG Interpretation (3ª ed.). Demos Medical Publishing.

[47] Hidalgo VM, Letelier JC, Díaz J. The amplitude modulation pattern of Gaussian noise is a fingerprint of Gaussianity. arXiv[preprint]. 2022. arXiv:2203.16253. doi:10.48550/arXiv.2203.16253.

[48] Charlebois, C. M., Anderson, D. N., Smith, E. H., Davis, T. S., Newman, B. J., Peters, A. Y., Arain, A. M., Dorval, A. D., Rolston, J. D., & Butson, C. R. (2024). Circadian changes in aperiodic activity are correlated with seizure reduction in patients with mesial temporal lobe epilepsy treated with responsive neurostimulation. Epilepsia, 65(5), 1360–1373. 10.1111/epi.17938

[49] Park, I., Díaz, J., Matsumoto, S., Iwayama, K., Nabekura, Y., Ogata, H., Kayaba, M., Aoyagi, A., Yajima, K., Satoh, M., Tokuyama, K., & Vogt, K. E. (2021). Exercise improves the quality of slow-wave sleep by increasing slow-wave stability. Scientific Reports, 11, 4410. 10.1038/s41598-021-83817-6

[50] Park, I., Kokudo, C., Seol, J., Ishihara, A., Zhang, S., Uchizawa, A., Osumi, H., Miyamoto, R., Horie, K., Suzuki, C., Suzuki, Y., Okura, T., Díaz, J., Vogt, K. E., & Tokuyama, K. (2022). Instability of non-REM sleep in older women evaluated by sleep-stage transition and envelope analyses. Frontiers in Aging Neuroscience, 14, 1050648. 10.3389/fnagi.2022.1050648

