## Supplementary Data for "Envelope analysis as a tool for identifying epileptic EEG patterns during the sleep-wake cycle in rats"

**Supplementary Materials**

**Supplementary 1:** Electrodes location for polysomnographic recording

| **EEG Electrode** | **Antero-Posterior** | **Lateral** |
| --- | --- | --- |
| 1 | 5.6 | -4.5 |
| 2 | 5.6 | 4.5 |
| 3 | - | -.5 |
| 4 | -2 | 1.5 |

Stereotactic coordinates for cortical electrode placement. For more detail review the scheme below.

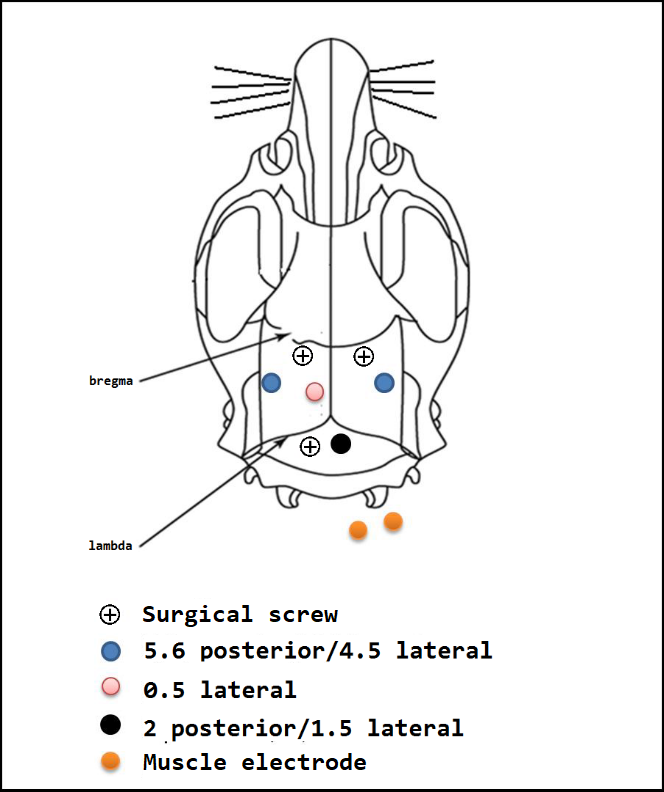

Electrode location scheme for polysomnographic recording. Cortical electrodes 1, 2 (blue), 3 (pink), and 4 (black) are identified. Electrodes for recording muscle activity are indicated with orange dots.

**Supplementary 2:** Anesthetic, antibiotic, and inflammatory management protocols.

**Anesthetic protocol**

| *Induction* | Xylacine 0.1 mg/Kg. + Ketamine 0.1 mg/Kg. |
| --- | --- |
| *Dose* | 0,15 ml Xylacine + 0,15 ml Ketamine dissolved in 0.7 ml of 0.9% physiological solution. |
| *Administration* | Intraperitoneal injection. |
| *Administration interval* | At the beginning of the surgical stage, once. |
| *Anesthetic conduction:* | Ketamine 0.1 mg/Kg. |
| *Dose* | 0,20 ml Ketamine dissolved in 0,8 ml of 0,9% physiological solution. |
| *Administration* | Intraperitoneal injection. |
| *Administration interval* | During the surgical procedure, if the animal shows signs of pain, with a maximum of 3 times. |

**Antibiotic protocol and management of the inflammatory process**

| *Antibiotic* | Enrofloxacin 10% |
| --- | --- |
| *Dose* | 0,1 ml dissolved in 0,4 ml of 0,9% physiological solution |
| *Administration* | Intraperitoneal injection |
| *Anti-inflammatory* | Ketoprofen 1% |
| *Dose* | 0,1 ml dissolved in 0,4 ml of 0,9% physiological solution. |
| *Administration* | Intraperitoneal injection |

The indicated doses were placed at the end of the surgical procedure. All procedures were recorded in the surgical record.

**Supplementary 3: Delta Envelope Characterization Space for one day of recording per rat.**

Scatterplot represents the amplitude of the delta band (y-axis) and coefficient of variation of the envelope (x-axis), and vertical dashed lines represent theoretical values for Gaussianity according to the sampling distribution of delta-CVE. Each rat describes the triangle-shaped distribution which is not observed in the model established by Diaz et al. [16] for wild-type.

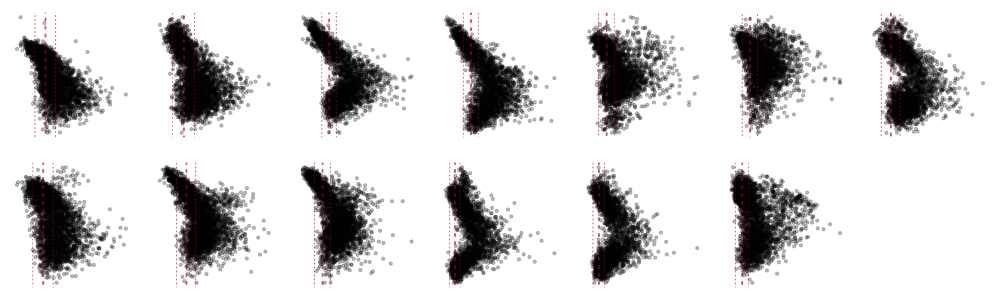

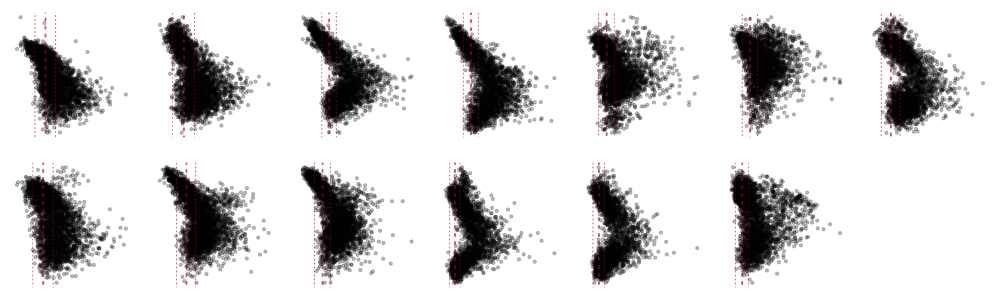

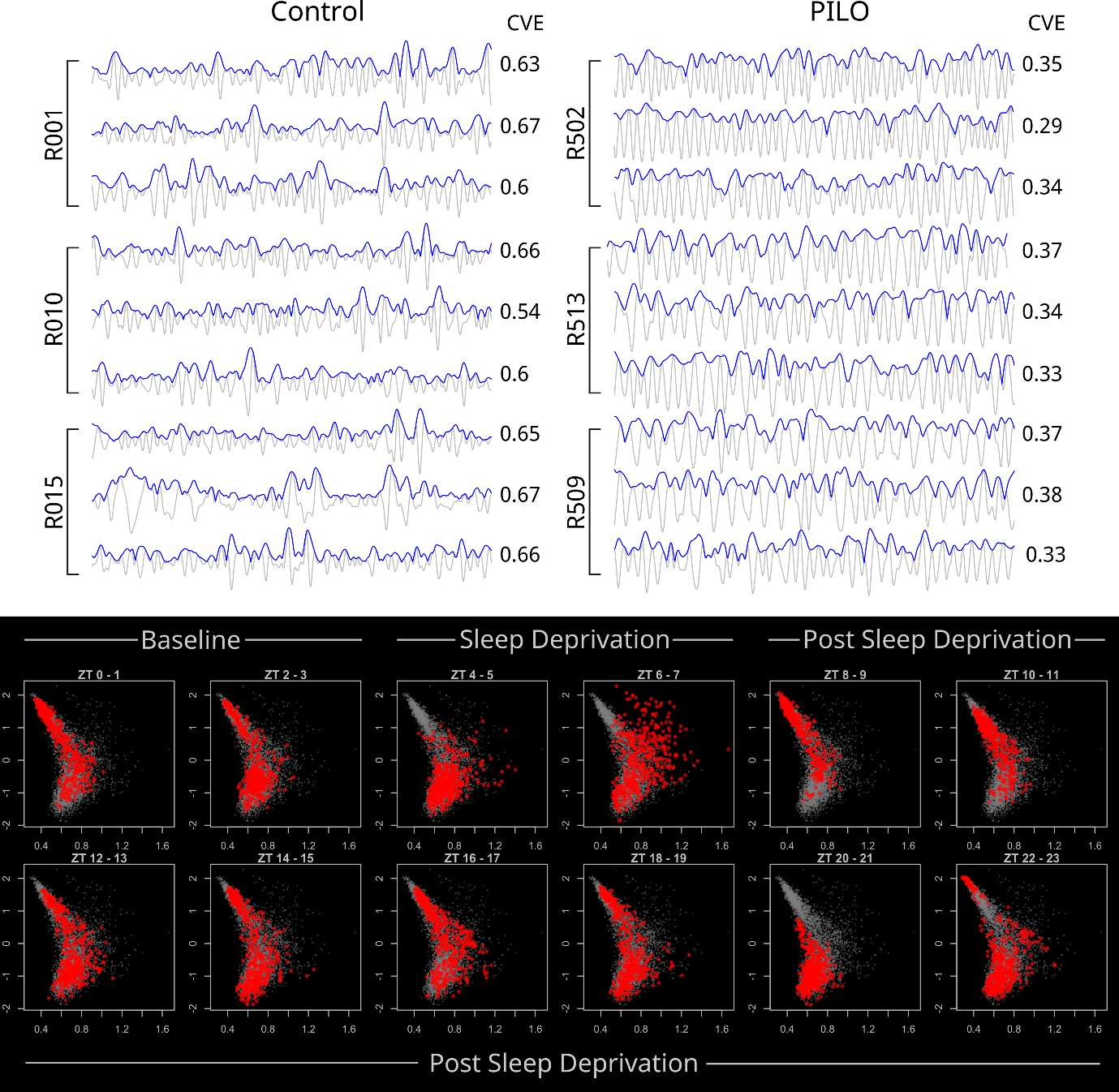
**Supplementary 4. Envelope analysis for delta activity in representative epochs per group, and Sleep deprivation Envelope Characterization Space.**

Envelope Characterization Space (ECS) for the sleep deprivation protocol. Each block indicates a stage of the protocol from left to right: baseline (ZT 0-3), sleep deprivation (ZT 4-7), and post-sleep deprivation (ZT 8-23). Gray dots represent all epochs, while red dots indicate epochs at each ZT. ZT: zeitgeber time; time after switching on the light. After sleep deprivation period, ZT4-5 and ZT6-7, is it possible that all dots reach the Q1 in ZT8-9, which suggests a homeostatic rebound, like in canonical non-REM sleep. This is evidence that sDelta, is in fact an altered non-REM sleep.
